# Emergence of Orientation Pinwheels in a Self-Evolving Spiking Neural Network: Enhancing Visual Coding Efficiency and Reliability

**DOI:** 10.1101/2024.03.08.583887

**Authors:** Haixin Zhong, Haoyu Wang, Mingyi Huang, Wei P Dai, Yuchao Huang, Mengyao An, Anna Wang Roe, Yuguo Yu

## Abstract

Orientation preference maps (OPMs) in the primary visual cortex of primates organize orientation-tuned neurons into columnar structures, forming pinwheel-like patterns. However, lower-level animals like rodents typically exhibit a lack of OPMs, with neurons either randomly distributed or aggregated in small clusters. This distinction prompts an inquiry into whether more structured cortical columns correlate with improved visual computational or coding efficiency. To explore this, we propose a novel self-evolving spiking neural network (SESNN). To the best of our knowledge, the SESNN is the first spiking network, incorporating mechanisms of neural plasticity in forming neural connections without explicit objective functions. We reveal that the emergence of pinwheel structures is primarily driven by sparse coding constraints and local synaptic plasticity as fundamental mechanisms. Second, for higher mammals with expansive iso-orientation domains (IODs), the firing responses in pinwheel structures primarily emanate from pinwheel centers (PCs) and progressively extend toward the periphery, encompassing adjacent IODs. Third, the size and organization of these IODs across species are significantly influenced by the receptive fields’ ability to process overlapping visual information. Lastly, PCs within large IODs demonstrate enhanced robustness and population sparseness in detecting a variety of orientation features. These results indicate that the spatial pinwheel structure facilitates highly efficient and reliable coding performance.

## 1. Introduction

Previous research has demonstrated that neurons in the primary visual cortex (V1) develop orientation selectivity through the sparse coding principle, enhancing visual neural representation and aiding in the creation of brain-inspired algorithms(Hubel & Wiesel, 1962; Vinje & Gallant, 2000; Bell & Sejnowski, 1997; Ba & Dj, 1997; Hubel, 1959; Field, 1987). In numerous species, V1 neurons are organized in pinwheel structures for orientation, which shift in orientation preference (Li et al., 2019; Bock et al., 2011; Bonhoeffer & Grinvald, 1991; Obermayer et al., 1992). Nonetheless, there is still a lack of understanding regarding how neurons within iso-orientation domains (IODs) and within pinwheel centers (PCs) of pinwheel structures across different species represent visual targets efficiently in the external environment.

Over the last forty years, the self-organizing map (SOM) algorithm (Kohonen, 1982) has played a pivotal role in reproducing OPMs at the mesoscopic level. Despite its contributions, the SOM falls short in capturing neural dynamics on both microscopic (single-neuron) and macroscopic (population-neuron) scales, thus limiting its utility in elucidating the functional significance of OPMs. To bridge this gap, we propose the self-evolving spiking neural network (SESNN) model, a cutting-edge 2D cortical framework tailored for OPM development mechanism.

Consistent with experimental evidence (Holmgren et al., 2003; Hofer et al., 2011), V1 pyramidal neurons exhibit weaker synaptic strengths compared to other types, crucial for avoiding over-excitation and ensuring neural balance. To accurately model these self-evolving dynamics within our SESNN, we implement excitatory-to-excitatory (E-E) connections governed by the Hebbian Oja’s rule (HO) (Oja, 1982), with a normalization factor (the second term in equation (4)) to regulate synaptic weight growth within a 0 to 1 range. In contrast, lateral connections between excitatory and inhibitory neurons under the correlation measuring (CM) rule are inherently stronger without such a normalization factor (Zylberberg et al., 2011; Paul D. King et al., 2013). In the 2D cortical model for neural connectivity, initial weights are determined by a Gaussian function reflecting neuron distance(Tao et al., 2004; Stepanyants et al., 2009; Amatrudo et al., 2012), incorporating a periodic boundary condition (PBC) to emulate V1’s complex network. This setup, coupled with retinotopic data on overlapping receptive fields of adjacent excitatory neurons, enhances the model’s realism(Srinivasan et al., 2015; Tehovnik & Slocum, 2007; Scholl et al., 2013; Veit et al., 2014; Engelmann & Peichl, 1996; Weigand et al., 2017; Law et al., 1988; Huberman et al., 2006; Niell & Stryker, 2008; van Beest et al., 2021; Foik et al., 2020).

### Our contributions are threefold

- Our SESNN model represents an unprecedented approach to replicate the spiking mechanism in orientation pinwheel structures across species. It uniquely generates sparse codes from natural scene learning via local synaptic plasticity, setting a new standard for neural coding strategy research.
- Our SESNN model aligns with empirical data, underscoring the role of visual input overlap in cortical architecture. It unveils a novel early-response mechanism where visual neurons in pinwheel structures exhibit unique firing patterns upon external stimuli, with center neurons firing first. This pattern, absent in lower animals, indicates pinwheel structures may use precise spiking timings for efficient coding, marking a significant discovery in neural coding strategies.
- Our findings reveal that higher mammals display enhanced sparse coding and noise resilience in their organized pinwheel structures, unlike lower mammals like rodents. PCs in those advanced mammals are notably more robust than non-pinwheel area, crucial for accurate orientation detection amidst noise. This highlights the evolutionary advantage of organized pin-wheel structures in complex neural processing.

## 2. Methods

### 2.1. Realistic Self-Evolving Spiking Neural Network

In this paper, we propose the SESNN model stimulated by natural images and governed by local synaptic plasticity rules, which are Hebbian-Oja (HO) and Correlation Measuring (CM) (Paul D. King et al., 2013; Zylberberg et al., 2011). The core component of our framework is depicted in Figure 1. To represent efficiently the input natural images, the SESNN for high-level animals self-evolves pinwheel structures, constrained by sparse coding algorithms. We aim to explore the self-organization processes within the primary visual cortex across various species in response to natural image stimuli and comprehend the resulting neural representations. The forthcoming sections provide a detailed description of the key elements of our model. These components encompass the degree of visual overlap in input stimuli reception, the architecture and connectivity of the model, and the specific rules of plasticity that guide learning.

**Figure 1.**
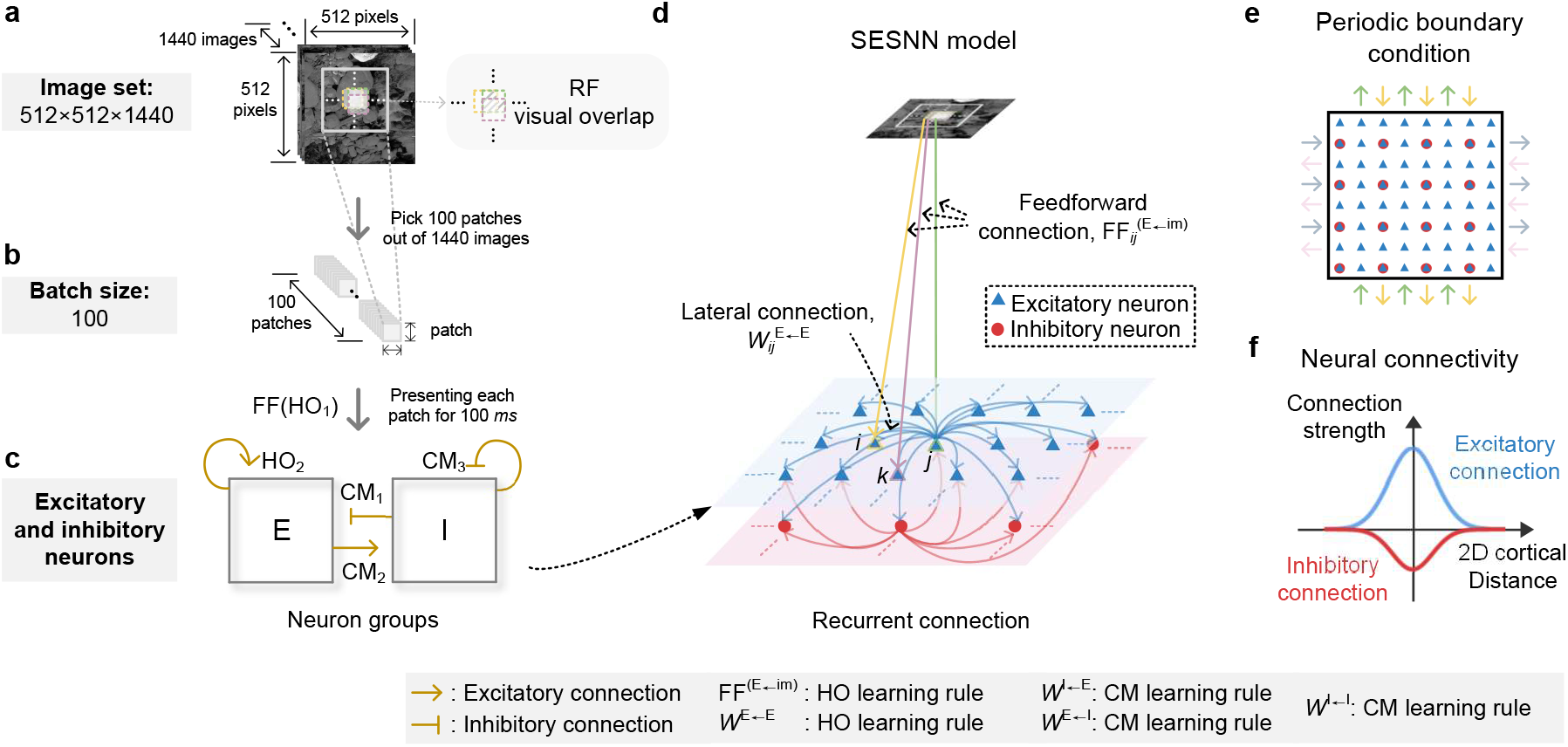
Architecture of proposed SESNN model. **a**. The image set contains 1440 natural whitening images of 512×512 pixels each. E-neurons have overlapping visual fields, each perceiving a 16×16 pixel patch. We use three levels of overlapping visual fields of three nearby E-neurons as an illustration. **b**. Stimulus images of 100 patches are picked out from the image set randomly and presented to E-neurons for 100 *ms* (Feedforward connection, following HO learning rule). **c**. The connections between E- and I-neurons (4900 and 1225) follow HO and CM rules. **d**. Recurrent connection among neurons. The feedforward connection FF^(*E←*image)^ follows the HO learning rule. All remaining lateral connections, *W* ^(*I←E*)^, *W* ^(*E←I*)^, and *W* ^(*I←I*)^, follow the CM learning rule. **e**. The spatial arrangement and PBC of E- and I-neurons. E- and I-neurons share the same spatial coordinates, and the boundaries are linked together according to the arrows shown in the diagram. Arrows of the same color represent identical connections. **f**. The initial weight distribution exhibits a Gaussian function. The values of initial weights are described by Equation 3, determined by the minimum distance between neurons considering the PBC described in Figure 1e.

#### 2.1.1. Stimulus training sets

The selection of stimulus inputs is pivotal in determining the distribution of orientation-preferred neurons within the emergent OPMs following training in SESNN. To this end, we use 20 original whitened images (512×512 pixels) which eliminate pairwise correlations among pixels, emulating the processing of the lateral geniculate nucleus (LGN) (Zylberberg et al., 2011). To ensure these images encompassed orientation details, each image is rotated clockwise every 5 degrees, yielding 72 images per original image. This process results in a total of 1440 images comprising our training data set (refer to Figure 1a).

In each trial, every E-neuron perceives 100 different patches for 100ms, with each 16×16 patch arbitrarily selected from the training data set to serve as the neuron’s RF (refer to Figure 1(b)). We assume the visual fields of the E-neurons to be overlapping on the retina (refer to Figure 1a). The overlap degrees in visual fields vary across species (including macaques, cats, rodents, etc.) to accommodate the size of IODs in the visual cortex representation as observed by experimental studies (Najafian et al., 2022).

To accommodate the visual overlap among adjacent neurons, which adhere to PBC, the visual input is designed to exhibit a wraparound effect. This ensures continuity and uniformity in processing across the neural network.

#### 2.1.2. Neural dynamics

The SESNN model comprises 4900 E- and 1225 I-neurons connected recurrently with an E-I ratio of 4:1 (e.g., cat, macaque) and 3600 E- and 400 I-neurons with 9:1 (e.g., rodent)(Alreja et al., 2022; Tao et al., 2004; Pfeffer et al., 2013). E-neurons receive stimuli from natural images as well as background noise from other brain areas. I-neurons indirectly receive natural image stimuli by adjusting E-neurons. The neural spiking dynamics are modeled using biologically-inspired leaky integrate-and-fire (LIF) neurons (Stevens et al., 2013), incorporating refractory periods (3 ms) and adaptive firing thresholds (Földiák, 1990). The neural dynamics are iteratively formulated as follows:

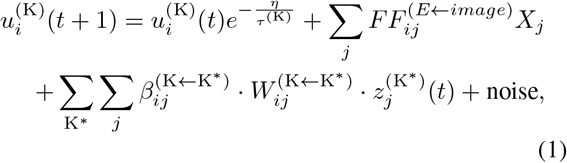

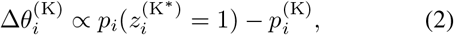

where K = E-neurons or I-neurons; *i* = 1, 2, …, *N* (numbers of E-neurons and I-neurons).

In Equation 1, 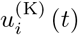 represents the membrane potential of neuron *i* at time *t* (subjects to neurons of class K). The resistor-capacitor circuit membrane time constant, denoted by *τ*, controls the decay rate for the membrane potential in individual neurons. We set *τ* ^(*E*)^ = 10 *ms* for E-neurons and *τ* ^(*I*)^ = 5 *ms* for I-neurons. It is worth mentioning that I-neurons are designed to fire at a faster rate than E-neurons (Paul D. King et al., 2013; Thomson & Lamy, 2007). This configuration facilitates decreased reconstruction error and accelerates the convergence of the system, thereby contributing to a more efficient and precise representation of the input stimuli. The feedforward (FF) connection, de-noted as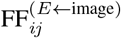, which links pixel *X*_*j*_ of the whitened image patch (in which pixels has been normalized to zero mean and unit variance) to E-neuron *i*. 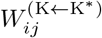 represents the connection weight from neuron *j* of neuron class K^*∗*^ to neuron *i* of neuron class K with its sign determined by the connection type, described as 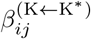 (the neuron receives excitatory connections, set as +1; conversely, the neuron receives inhibitory connections, the sign is set as -1). 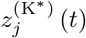 refers to the spike output of neuron *j* at time *t*. After reaching the spike threshold *θ* (initialized as 2), a spike is emitted, 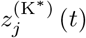 is set to 1, then the membrane potential is reset to 0 mV and holds until the end of the refractory period. The firing threshold *θ* continuously adapts based on the difference between the current firing rate *p*_*i*_ (*t*) and target firing rate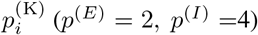 within a 10 ms time window, as depicted in Equation 2(Földiák, 1990). For computational efficiency, we set the time step *η* to 1 ms.

#### 2.1.3. Neural connectivity within 2d cortical area

In our SE-SNN model, E- and I-neurons are arranged symmetrically on a two-dimensional lattice, as illustrated in Figure 1e. PBC is employed to mimic the large number of neurons in the actual V1. Specifically, neurons at the boundary are connected to neurons at corresponding symmetric positions on the opposite boundary. The initial connection weights between neurons are modeled by a Gaussian function of physical distance, which can be expressed as:

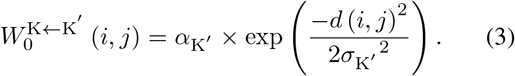

In this equation, *d*(*i, j*) represents the Euclidean distance from neuron *i* to neuron *j* in a grid, *α* determines the maximum connection weight, which is set to *α*_EE_ = 1, *α*_EI_ = 1, *α*_IE_ = 0.5, *α*_II_ = 0.5, and *σ* governs the rate at which the weight decays with distance. The synaptic types predominantly determine the parameters for this connection weight distribution function. To accurately replicate the neuronal architecture of V1 in macaques. The connectivity radiuses, denoted by *σ*, are set to *σ*_EE_ = 3.5, *σ*_EI_ = 2.9, *σ*_IE_ = 2.6, *σ*_II_ = 2.1. These values are based on anatomical data indicating that the axon lengths of E- and I-neurons are approximately 200 *µ*m and 100 *µ*m, respectively, while the dendrite lengths are around 150 *µ*m for E-neurons and 75 *µ*m for I-neurons in the V1 (Tao et al., 2004; Stepanyants et al., 2009; Amatrudo et al., 2012). This careful selection of parameters and neuron distribution ensures our model’s biological fidelity, particularly in representing synaptic connectivity and neuronal density. We discard any connection strengths below a threshold of 0.01 to maintain computational efficiency and biological plausibility.

#### 2.1.4. neural plasticity rules

The learning process is remarkable because it does not rely on external teaching signals. Two forms of Hebbian learning rules primarily govern self-learning processes: the Hebbian Oja’s variant (Oja, 1982) and the correlation measuring rules (Zylberberg et al., 2011; Paul D. King et al., 2013). These rules facilitate the model’s ability to adaptively adjust synaptic weights based on the correlation of firing patterns, thereby mimicking a fundamental aspect of learning in biological neural networks. As introduced in the Introduction 1 section, we manage E-E and FF connections with the HO synaptic rule, while others with the CM rule. Both HO and CM rules have analogous mechanisms that implement long-term potentiation (LTP) and depression (LTD), like most other rate learning rules that impose no constraint on the exact timing of spikes, thus the learning rules can be relaxed. The reason that we choose HO and CM rules is because less tuning efforts are needed, and stability in recurrent excitation.

Following every 100 ms of stimulus presentation, synaptic weight changes within the network are computed. The formula for these adjustments is given by:

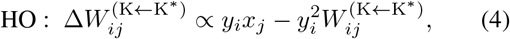

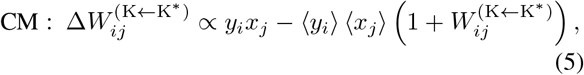

where *x, y* refer to the spike rate of presynaptic and postsynaptic neurons, the operator ⟨·⟩ denotes the lifetime average value. Hyperparameters of the SESNN model are presented in Table 1.

**Table 1.** SESNN model hyperparameters.

| PARAMETER | VALUE | DESCRIPTION |
| --- | --- | --- |
| **LEARNING RATE** |  |  |
| $\eta_{FF}$ | 0.2 | IMAGE TO E-NEURON |
| $\eta_{EE}$ | 0.01 | E- TO E-NEURON |
| $\eta_{EI}$ | 0.7 | I- TO E-NEURON |
| $\eta_{II}$ | 1.5 | I- TO I-NEURON |
| $\eta_{IE}$ | 0.7 | E- TO I-NEURON |
| **NEURAL CONNECTIVITY** |  |  |
| $\alpha_{\text{MAX},E}$ | 1.0 | E- MAX WEIGHT |
| $\alpha_{\text{MAX},I}$ | 0.5 | I- MAX WEIGHT |
| $\sigma_{EE}$ | 3.5 | E-E COUPLING RANGE |
| $\sigma_{EI}$ | 2.9 | E-I COUPLING RANGE |
| $\sigma_{IE}$ | 2.6 | I-E COUPLING RANGE |
| $\sigma_{II}$ | 2.1 | I-I COUPLING RANGE |

**Table 2.** SESNN pinwheels vs. macaque pinwheels (mean ± SD).

| METRIC | SESNN | MACAQUE |
| --- | --- | --- |
| PINWHEEL DENSITY (PINWHEELS/ $\Lambda^2$) | $3.175 \pm 0.397$ | $\sim 2.934$ |
| NEIGHBOR DISTANCE (MM) | $0.277 \pm 0.043$ | $\sim 0.209$ |
| HYPERCOLUMN SIZE (MM) | $0.839 \pm 0.054$ | $\sim 0.657$ |

### 2.2. Anatomical Data Integration

The anatomical data for neural connectivity in our study is based on the findings of (Tao et al., 2004; Stepanyants et al., 2009; Amatrudo et al., 2012). The plasticity rule settings are based on the findings of (Holmgren et al., 2003; Hofer et al., 2011). For retinotopic data, we refer to (Srinivasan et al., 2015; Tehovnik & Slocum, 2007; Scholl et al., 2013; Veit et al., 2014; Engelmann & Peichl, 1996; Weigand et al., 2017; Law et al., 1988; Huberman et al., 2006; Niell & Stryker, 2008; van Beest et al., 2021; Foik et al., 2020). The anatomical data on OPMs is sourced from (Najafian et al., 2022).

## 3. Results

### 3.1. Visual overlap underlying orientation map formation across species

Our proposed SESNN model emerges the orientation map successfully (refer to Figure 2f-g). Simultaneously, the visual overlap among RFs, which perceive overlapping natural images, is a principal determinant in differentiating species. According to our results in Figure 2a, this overlap is a critical element in the development of the diverse IODs across species, such as rodents, macaques, and cats. In our model, we adjust the overlap parameters, ranging from 9 to 15 pixels, to increase the size of the IODs. Notably, when the overlap is set at 9 pixels, the model generated a salt- and-pepper organization instead of the expected pinwheel structure, as depicted in Figure 2a. Consequently, when the overlap is equal to 9 pixels, we exclude it from the analysis of the pinwheel structure, as reflected in Figure 2b-d.

**Figure 2.**
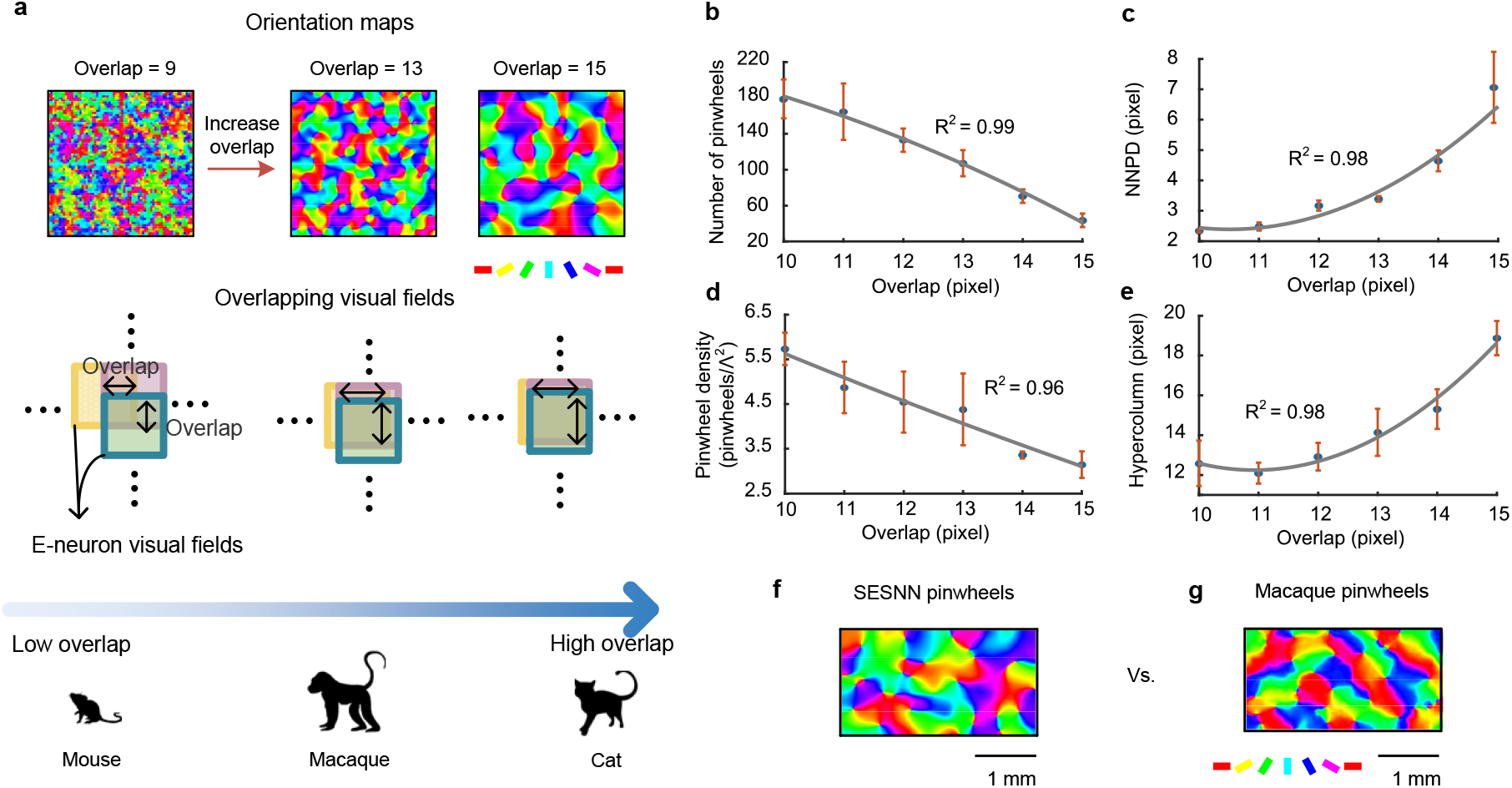
OPMs emergence and neuronal responses via our SESNN model. **a**. The overlap of visual fields in neurons, perceiving overlapping natural images, forms various IODs of species in the OPMs. The increase in the overlap is consistent with the transition from lower to higher mammals (see the animal sketches). **b**. The correlation between visual overlap and mean PC counts (± SD). **c**. The relationship between visual overlap and NNPD (± SD). **d** Variability in mean pinwheel density (± SD) with different visual overlap. **e**. The correlation between visual overlap and mean hypercolumn size (± SD). **f**. SESNN pinwheels vs. Macaque pinwheels. The scale bar corresponds to 1 mm on the V1 cortical surface. The color bars represent orientation preferences.

To quantitatively analyze the orientation maps presented in Figure 2a, several measures are employed (Najafian et al., 2022), including the number of PCs, nearest neighbor pin-wheel distance (NNPD), PC density, and size of the hyper-column. These measures are illustrated in Figure 2b. Figure 2c describes the relationship between the NNPD and the overlap among RFs. The NNPD is defined as the distance between the two nearest PCs. Figure 2d shows the relationship between the number of PCs and overlapping visual fields. PCs can be measured by 2D FFT (Kaschube et al., 2010), which are located at the intersection of the real and imaginary components that equal 0 (Stevens et al., 2013). Pinwheel density can be calculated as the number of PCs in a hypercolumn (Λ^2^), as shown in Figure 2d. This panel also demonstrates how overlapping visual fields relate to pinwheel density. Moreover, we defined a hypercolumn as a region with periodicity measured by 2D FFT. Figure 2e shows how overlapping visual fields relate to the size of the hypercolumn.

Additionally, to substantiate the SESNN model’s results, we introduce biological data from macaque pinwheels for comparison (see Figure 2f-g and Table 1).

### 3.2. Spatial-temporal distributed spiking waves propagate within pinwheels

Figure 3 illustrates our findings on higher mammals with large IODs, such as macaques and cats. A quantitative depiction of the findings in Figure 3a is presented in Figure 3b. Our study reveals that V1 neurons, which are stimulated by natural images, primarily fire within the pinwheel orientation structures originating from the PCs or IODs located near the PCs and extending towards the periphery. This phenomenon is particularly notable in species with large IODs.

**Figure 3.**
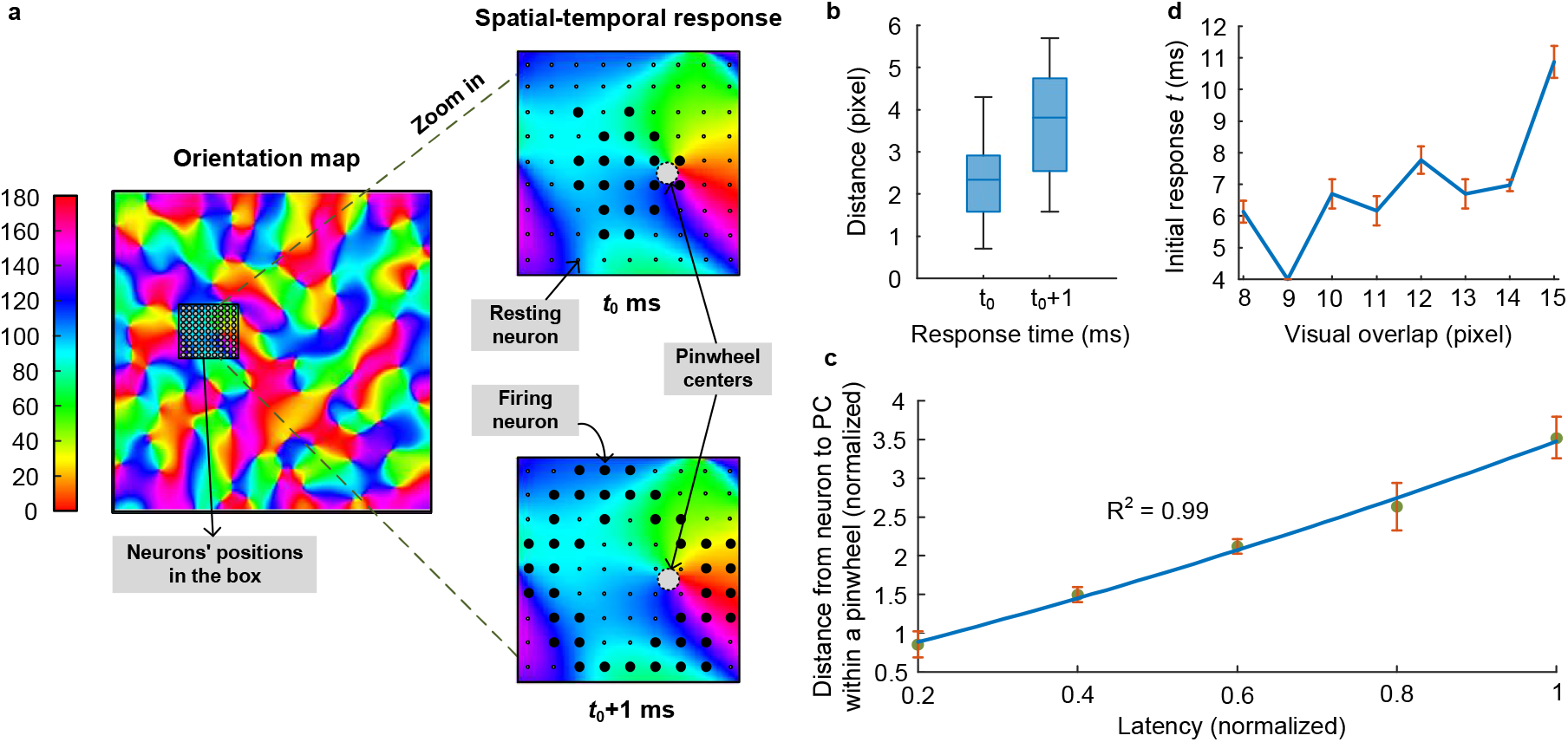
Spatial-temporal response structure within pinwheels. **a**.This figure displays the neuronal responses on OPM with a large IOD in a pinwheel structure. The neurons that fire at time *t*_0_ are shown as large black dots at PC, and they expand towards the periphery at time *t*_0_+1, also denoted as large black dots. The other small dots represent resting neurons. **b**. Distance between firing neurons and the PC at time *t*_0_ and *t*_0_+1. **c**. This panel shows the latency of neurons and the mean distance (± SD) between these neurons within a pinwheel. **d**. The figure illustrates the mean values of initial response time (± SD) in different species with varying sizes of IODs, ranging from small to large IODs.

In Figure 3c, We select a representative example where neuronal responses originate from the center of a pinwheel structure and conduct an analysis recording the timing of responses and their distances from the PC. We define the onset of discharge from the center as a latency of 1 ms, with subsequent firings within the next time step having a latency of 2 ms, based on our 1 ms time unit. Under natural external image stimulation, we observe discharges starting at the center and exhibiting pronounced diffusion. Neurons within the pinwheel IODs activate sequentially, depending on their distance from the center, showing a clear diffusion pattern.

We then conduct an extensive investigation into the SESNN model, subjecting it to continuous stimulation with varying degrees of overlapping receptive fields. To achieve this, we utilize the noise-free natual image, as illustrated in Figure 4a, as the external stimulus. The primary objective is to analyze the precise timing at which E-neurons within the model exhibit their initial firing activity. By performing 30 experimental trials, we obtain the mean and variance of the initial firing times, as depicted in Figure 3d. Our results demonstrate a significant increase in the average initial response time as the degree of overlapping receptive fields increases. This finding indicates that higher animals exhibit a longer response time compared to lower animals. Remarkably, this observation aligns coherently with our conclusions regarding the sparsity characteristics of the SESNN model (see below). As the complexity of the species increases, the model’s capacity to represent and process information improves, resulting in a higher level of sparsity in the model’s responses. Consequently, the temporal responsiveness of neurons to stimuli becomes delayed following stimulation.

**Figure 4.**
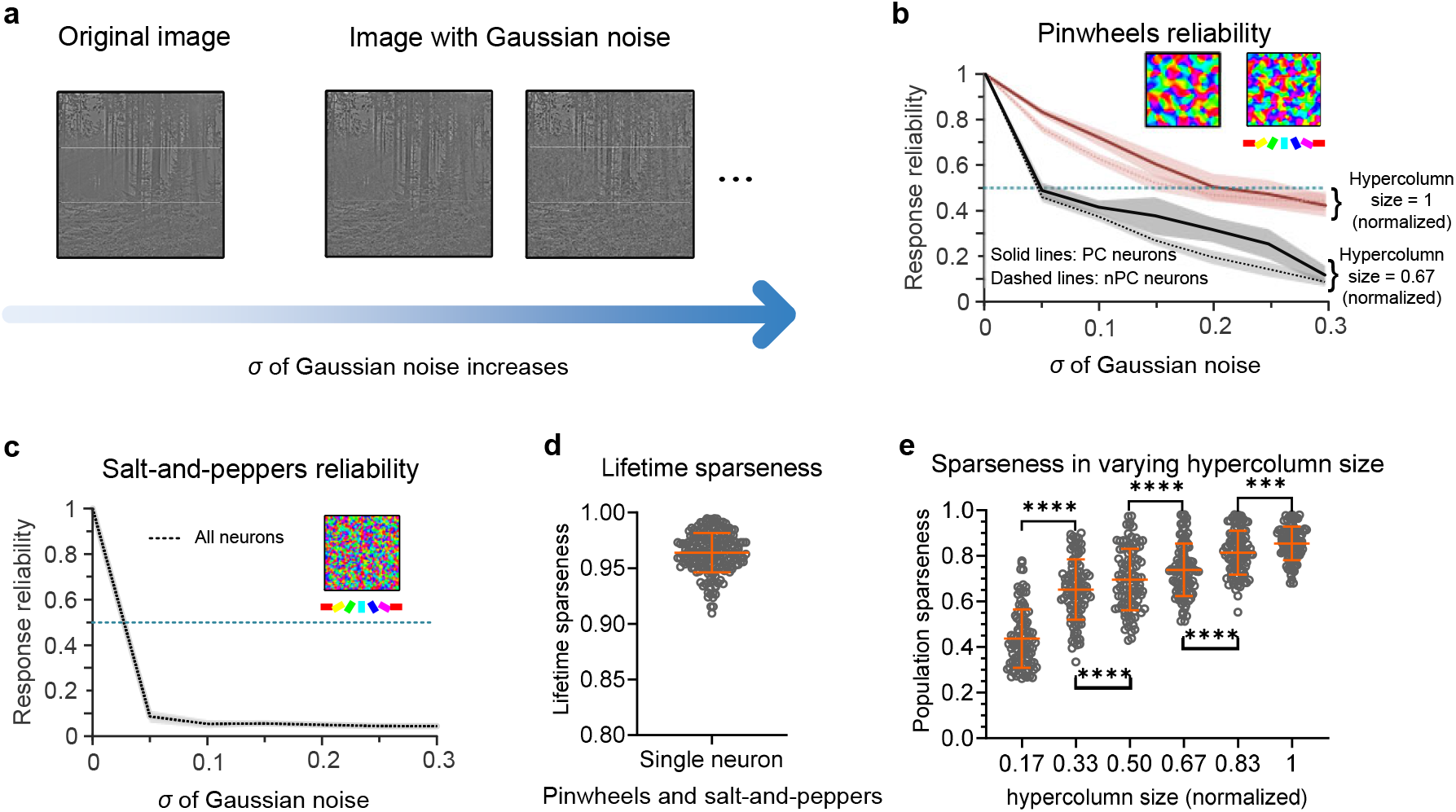
Populational responses of pinwheels and salt-and-peppers. **a**. The left panel displays image inputs with different levels of Gaussian noise (*σ* ranges from 0 to 0.3) applied to our trained SESNN model. **b-c**. Response reliability of both pinwheels (different hypercolumn sizes) and salt-and-peppers. Insets visualize these patterns, with color bars indicating orientation preferences. The dashed blue line y = 0.5 is a benchmark to facilitate the comparison of different neuron types at a particular level of noise. **d**. Lifetime sparseness of each neurons across species. **e**. Population sparseness of pinwheels across different hypercolumn sizes. (All data: mean ± SD, ^***^ p*<*0.001, ^****^ p*<*0.0001).

### 3.3. Populational response of pinwheels and salt-and-peppers

Now, we investigate the response reliability of neurons under the stochastic condition, which is crucial for trustworthy and robust neural information processing. To quantify response reliability, we employ a spike-based measurement approach. For each neuron, we analyze spike time sequences produced under identical stimulus conditions with added noise, calculating the mean cross-correlation between these sequences. We then determine the maximum average cross-correlation within a maximum delay time not exceeding half the length of the time sequence as the neuron’s reliability. For each neuron, *n* spike trains of duration time *T* are analyzed. After normalizing the mean of each train to zero, we systematically calculate the cross-correlation coefficient between each pair of sequences from these *n* sequences, employing the following approach:

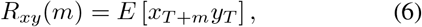

where *m* denotes lag, the range of which can extend over the length of the time sequence *T. x* and *y* denote two different spike trains. Through this procedure, the average correlation coefficient of the same neuron can be determined:

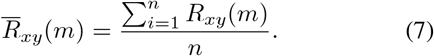

We posit that the maximum lag for similar time sequences generated by the same neuron cannot exceed half the length of the time sequence. Consequently, the reliability is obtained as follows:

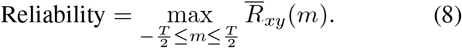

To evaluate the SESNN model’s response reliability to noise-corrupted natural images as stimuli, we introduce Gaussian random noise (mean of 0, variance up to 0.3) to the images. The model is stimulated with this variably noisy image ten times. We calculate the response reliability for neurons at both PCs and nPCs. However, given the SESNN model’s 9-pixel overlapping receptive fields do not efficiently produce pinwheel structures, we do not distinguish between neuron types. The reliability values obtained are depicted in Figure 4b-c.

We predict theoretically the phenomenon for the first time that as the IOD sizes increase from small to large across different species (refer to Figure 4b), the response reliability of neurons within the pinwheels also increases. This enhanced reliability allows these neurons to maintain stable response activities even in the presence of noise interference, highlighting the superiority of higher mammals. Furthermore, within the same species, we observe that neurons located at the PC neurons exhibit higher reliability compared to nPC neurons. Subsequent experimental studies could validate these theoretical forecasts.

In Figure 4e, we calculate the population sparseness (Willmore et al., 2011) and lifetime sparseness (Rolls & Tovee, 1995) in various species when presented with 100 natural image patches of a 100 ms time window each. Lifetime sparseness is defined as the distribution of a single neuron’s responses across multiple stimuli, defined as:

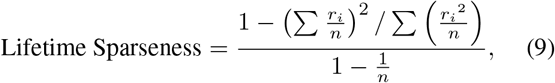

where *r*_*i*_ refers to spikes of a neuron triggered by the *i*-th picture out of a total of *n* pictures perceived by the neuron’s RF. Population sparseness, on the other hand, refers to the distribution of responses among a population of neurons to a single patch, as depicted below:

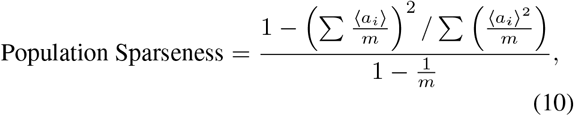

where *a*_*i*_ refers to the mean spike rate of a neuron *i* out of a total of *m* neurons in a time window.

The model results indicate that as the IDO sizes increase from small to large across different species, there is a tendency for population sparseness to increase. This is illustrated in Figure 4b, which displays scatter plots and columns representing the population sparseness in 100 patches, along with their median values. Additionally, Figure 4d presents the lifetime sparseness of different species, indicating each neuron emerges with specific selectivity.

## 4. Discussion

SOM has been utilized for decades to explain the emergence of orientation pinwheels (Oja, 1982). However, these models have notable limitations in predicting the spatiotemporal neuronal activities within visual circuits and understanding the coding principles employed by neurons within the pinwheel structures. Therefore, solely relying on self-organization models to comprehend pinwheel activity across different species poses challenges.

To the best of our knowledge, our SESNN is the first spiking network that incorporates biology-inspired LIF neurons and local plasticity rules, achieving a more precise and realistic representation of neuronal responses as well as sparse coding mechanism underlying the emergence of pinwheel structures. Furthermore, we emphasize the importance of different RFs perceiving overlapping information from the external world, which contributes to the emergence of IOD sizes across species (refer to Figure 2), such as mouse, macaque, and cat.

Our modeling studies indicate that species with more organized pinwheel structures should exhibit higher coding robustness (Figure 4b-c) and population sparseness (Figure 4e) compared to rodents lacking organized pinwheel structures. Specifically, the PCs in higher mammals demonstrate higher robustness than nPCs (solid line vs. dashed line in the same color in Figure 4b). This suggests that higher mammals, including cats and macaques, with larger IODs exhibit enhanced reliability and population sparseness in detecting different orientation features. Thus, PCs with large IODs play a crucial role in ensuring robust detection of various orientation features and exhibit higher coding efficiency.

The model results also demonstrate novel observations that the neurons in PCs exhibit initial firing responses to natural image stimuli and diffuse across IODs (refer to Figure 3a. This phenomenon indicates that higher mammals with higher population sparseness demonstrate more effective spatial-temporal responses that increase the coding capacity compared to lower animals, which have lower population sparseness (refer to Figure 4e). Furthermore, a neuron exhibiting high lifetime sparseness (Figure 4d) indicates that it is finely tuned to respond to particular features within the stimulus environment.

Our study may have a potential limitation in that it primarily examines OPMs on a local scale. This focus allows us to delve into local spatiotemporal population dynamics but limits our ability to grasp the broader network dynamics that large-scale networks, which require modeling of extensive neuron populations and long-range connections, might exhibit. Addressing this limitation by expanding our model to encompass these long-range connections will be a direction for our future study.

The further study could also focus on investigating the relationship between the initial response of PCs (Figure 3a) and the generation of saliency maps, which has the potential to advance our understanding of bottom-up processing mechanisms in the visual system. By exploring how the initial responses of PCs contribute to the generation of saliency maps, we can uncover the fundamental processes involved in detecting and prioritizing visually salient features in the environment. This line of inquiry holds great promise for advancing our understanding of attention and perception mechanisms. By exploring the organization and functioning of the visual system’s bottom-up processing, we can uncover valuable insights into the computational principles that underlie these processes. Furthermore, this research has the potential to inform the development of brain-inspired algorithms, such as sparse coding strategy (Figure 4e), image recognition in noisy environments (Figure 4b), and optimizing the E/I ratio for enhanced visual representation. These innovations simulate and replicate the intricate workings of the visual system.

### Impact Statements

This paper aims to advance the field of computational neuroscience, specifically within the realm of understanding the structural and functional dynamics of the visual cortex. By focusing on the development and application of advanced computational models, the research contributes to deeper insights into how complex visual information is processed and represented in the brain. This work is pivotal for bridging theoretical models and experimental findings, offering a comprehensive framework for exploring neural coding mechanisms, network connectivity, and the emergence of functional architectures in sensory processing systems. Additionally, the implications of this research extend to the design of brain-inspired algorithms, enhancing artificial intelligence systems’ ability to emulate human visual processing capabilities.

